# Local cooperative interactions reshape the folding transition in a one-dimensional spin-glass model

**DOI:** 10.64898/2026.06.30.735452

**Authors:** Rohon Mitra, Biman Jana

## Abstract

Protein folding is the process by which a polypeptide chain organizes into its three-dimensional structure through a balance of stabilizing and destabilizing interactions encoded by the sequence. A central question in protein biophysics is how thermodynamic factors guide a polypeptide toward its native folded state despite the rugged energy landscape and the competing influence of nonnative interactions. In many biomolecular processes, cooperativity provides a mechanism by which multiple weak interactions act collectively to generate a robust response. In the context of protein folding, such cooperative effects may arise when the formation of one native contact enhances the stability or likelihood of nearby native contacts, thereby promoting collective organization toward the folded state. At the same time, folding is opposed by the much larger number of non-native interactions, whose heterogeneity can introduce frustration and destabilize folding even when the average native bias favors the folded phase. The interplay of these competing effects in determining foldability remains unclear in statistical-mechanical models. Here, we address this problem using a one-dimensional spin-glass model of protein folding with explicit shared-residue cooperative interactions encoded through wedge-based motifs. We show that modest cooperative bias can stabilize folding even where the noncooperative system remains unfolded, whereas non-native energetic fluctuation suppresses folding and shifts the transition to higher cooperative strengths. We further find that partial cooperative coverage is sufficient to lower the folding threshold. Therefore, the model provides a mean-field framework for incorporating cooperative interaction strength into the native one-dimensional model of protein folding and for describing how local cooperativity reshapes the folding transition.

## Introduction

Protein folding is an intricate and highly regulated biological process in which a linear chain of amino acids transforms into a specific three-dimensional structure that determines function [1–8]. The classical Anfinsen hypothesis states that the information required for this process is encoded in the amino-acid sequence itself, implying that the native structure can, in principle, emerge from sequence-dependent physical interactions under appropriate conditions [1–3]. At the same time, folding is not a trivial search through all possible conformations. As emphasized by Levinthal’s paradox, a random exploration of configurational space would require an unrealistically long time for a protein to reach its native state [9–13]. Nevertheless, proteins typically fold on biologically relevant timescales, indicating that the sequence guides the system along organized and energetically favorable routes rather than through unguided sampling. This behavior is naturally described within the framework of energy-landscape theory, in which folding is viewed as stochastic motion on a high-dimensional free-energy surface containing local minima, metastable intermediates, and multiple competing pathways, while remaining overall biased toward the native basin [14–24]. In this framework, the amino-acid sequence does not merely specify the native structure; it also encodes the pattern of interactions that shapes the underlying free-energy landscape, thereby biasing the folding dynamics toward kinetically accessible and thermodynamically favorable conformations [10, 16, 21, 25]. Experimental and theoretical studies have further shown that folding may proceed through intermediates, parallel pathways, hierarchical organization, or nucleation-like events depending on the protein and the landscape under consideration [8, 26–30]. Although recent advances in sequence-to-structure prediction, particularly through AlphaFold, have shown that amino-acid sequence contains sufficient information to determine native structure with striking accuracy, they do not by themselves explain the underlying folding mechanism. The thermodynamic and kinetic principles through which sequence-encoded interactions shape the free-energy landscape and guide a protein to its native state therefore remain a central problem in protein biophysics [16–18, 31].

Statistical-mechanical studies of protein folding have led to a range of complementary minimal models, each formulated to describe a particular physical aspect of the folding process. Native-biased and structure-based models, including Gō-type models, emphasize the role of native contacts and overall topology in guiding the folding process [25, 31–35]. Ising-like models, particularly the Wako– Saitô–Muñoz–Eaton model, offer a tractable way to study folding thermodynamics and free-energy landscapes based on native structural information [36–38]. Minimal lattice heteropolymer models have likewise shown how sequence heterogeneity, frustration, and contact formation can produce distinct folding pathways [39–41]. Extensions and related reduced descriptions have also been used to study broader folding scenarios, including multiple native states and landscape-guided association processes [42, 43]. Within this broader class of physics-based models, the spin-glass framework provides a foundational statistical-mechanical description of protein folding and was originally developed in the context of disordered magnetic systems. In its adaptation to protein folding by Bryngelson and Wolynes, the spin-glass model represents the protein as a disordered system with heterogeneous interactions, where the folding transition is governed by the competition between native stabilization and frustration [14–16]. The mean-field random-energy description of this framework captures the unfolded, folded, and glassy phases of the system and provides a minimal picture of rugged free-energy landscapes and folding transitions [14, 15].

Despite their differences, many statistical-mechanical models of protein folding place primary emphasis on native contacts and the stabilization they provide as the principal driving force toward the folded state [25, 31, 35, 44–46]. In such a picture, folding is governed primarily by the energetic bias toward native interactions, although the much larger number of non-native interactions can yield a comparable net contribution through their collective statistical weight [47–51]. This observation suggests that the stability of the folded state may not be governed solely by independent native-contact bias. Instead, native residue pairs may form an interconnected network of mutually reinforcing interactions, so that one favorable native contact enhances the stability or likelihood of others. This viewpoint is also consistent with studies emphasizing landscape roughness, internal friction, specific non-native traps, electrostatic modulation, and environment-dependent slowing or acceleration of folding [52–58]. Recent theoretical and experimental work supports the idea that native contacts tend to organize into mutually reinforcing cooperative networks. Analysis of unbiased all-atom folding trajectories by Wang et al. showed that native contacts form the most cooperative pairs in the unfolded ensemble, and that the largest network of mutually reinforcing contacts is composed predominantly of native interactions [46]. Complementary recent work further suggested that cooperative native-contact formation facilitates barrier crossing and promotes folding by enhancing collective native stabilization [59]. Together, these results support the idea that native contacts do not act independently, but rather as cooperative networks that help drive the system toward the folded state [46, 59].

Cooperativity is a fundamental organizing principle in many biological processes, where the effect of one local event is amplified by others to produce a collective response [60, 61]. A classic example is hemoglobin, in which oxygen binding at one site increases the affinity of the remaining sites, enabling efficient and switch-like oxygen uptake and release [62–69]. Similar cooperative effects are central to allosteric regulation, protein–ligand binding, molecular recognition, and the assembly of multicomponent biomolecular complexes, where collective interactions often determine whether a process proceeds efficiently toward its functional state [61, 70–72]. More broadly, cooperative or collective effects have also been explored in contexts ranging from orientational glassiness and hydrophobic polyelectrolyte transitions to hydrogen-bond networks, microbial and ecological cooperation, and other collective dynamical settings [73–78]. In this sense, cooperativity provides a general mechanism by which many individually weak interactions can act together to produce a robust physical or biological outcome [60]. To the best of our knowledge, within spin-glass-inspired statistical-mechanical descriptions of protein folding, analytically tractable models that explicitly encode cooperative effects at the residue-pair–residue-pair level remain largely unexplored.

In this work, we develop a minimal theoretical framework to examine how local cooperative interactions reshape the folding transition in a one-dimensional spin-glass model of protein folding. By introducing wedge-based shared-residue cooperative motifs within a mean-field description, we investigate how explicit local cooperative stabilization modifies the folding landscape and influences the folding transition. This framework enables us to study how cooperative bias, native–non-native energy deviation, non-native energetic fluctuation, and partial cooperative occupation of native wedges jointly control the onset and stability of the folded state. Our results show that even a modest cooperative bias can induce folding, whereas non-native energetic fluctuation strongly suppresses it, consistent with the idea that small but coordinated networks of native interactions can stabilize folding while competing non-native heterogeneity opposes the emergence of the native state.

## Model

We extend the one-dimensional spin-glass model of protein folding developed by Wolynes and coworkers by introducing an additional residue-pair–residue-pair interaction term in the Hamiltonian to account for cooperative stabilization. In this description, protein folding is governed by a rugged energy landscape containing multiple metastable minima, and the folding process corresponds to the exploration of this landscape en route to the native functional state. The total energy of a given protein configuration is decomposed into distinct contributions as follows: *ε*_*i*_(*α*_*i*_) denotes the energy of the *i*^th^ amino acid residue in its conformation *α*_*i*_; *J*_*i,i*+1_(*α*_*i*_, *α*_*i*+1_) represents the nearest-neighbor interaction between successive residues along the peptide backbone; and *K*_*i,j*_(*α*_*i*_, *α*_*j*_, *r*_*i*_, *r*_*j*_) accounts for long-range interactions between residues that are brought into spatial proximity in the tertiary structure. Here, *α*_*i*_ specifies the state of residue *i* at sequence position *r*_*i*_, and each residue is assumed to possess one native state together with approximately ten non-native states (*ν* ≈ 10), as originally proposed by Wolynes and co-workers. The total energy of a configuration is therefore written as

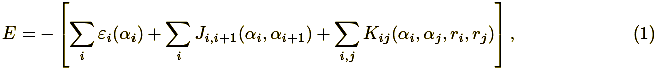

A direct analytical treatment of Eq. (1) is generally intractable owing to the exponentially large configurational space. We therefore adopt a stochastic Hamiltonian formulation in the spirit of Derrida’s random-energy model, wherein distinct conformations at fixed *N*_0_ are assumed to be statistically independent. Accordingly, for a chain of *N* residues with *N*_0_ residues in the native state, the joint probability distribution for *n* configurations with energies {*E*_*i*_} factorizes as 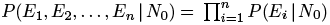. At the single-residue level, non-native conformational energies are assumed to be distributed with mean 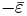 and standard deviation Δ*ε*, whereas the native-state energy is fixed at −*ε*_0_, with 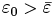. For nearest-neighbor interactions along the chain, non-native couplings are characterized by mean 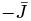 and standard deviation Δ*J*, while native couplings are assigned the fixed value −*J*, with 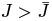. Similarly, at the tertiary level, non-native interactions are taken to have mean 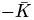 and standard deviation Δ*K*, whereas native tertiary interactions are fixed at *K*, with 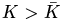. The average number of tertiary contacts per residue, denoted by *z*, is typically of order 2–3 in proteins, although it may vary weakly between the unfolded and folded states. Within the random-energy approximation, these one-body, nearest-neighbor, and tertiary contributions together imply that the total energy for configurations with *N*_0_ native residues is Gaussian distributed, with mean given by

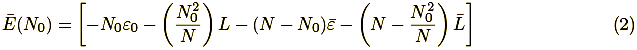

where

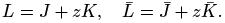

The corresponding standard deviation of the energy distribution, Δ*E*(*N*_0_), is

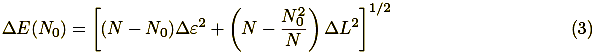

with

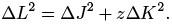

Equations (2) and (3) therefore constitute the baseline random-energy description of the folding model prior to the inclusion of any explicit cooperative stabilization energy.

### Shared-Residue (“Wedge”) Cooperativity

To incorporate local cooperative stabilization, we introduce an additional interaction associated with a *shared-residue contact pair*, or “wedge,” in which two contacts are anchored on the same central residue. In the present construction, we first consider the central residue to be native and then classify the wedge according to the states of the two partner residues connected to it. If both partner residues are native, then both incident contacts are native and the corresponding residue-pair–residue-pair motif is identified as a *native cooperative wedge*. By contrast, if only one of the two partner residues is native, or if both partner residues are non-native, then at least one of the two incident contacts is non-native; such motifs are therefore grouped together as *mixed cooperative wedges*. Thus, the native cooperative wedge corresponds to a fully native three-residue local environment, whereas all other possibilities are treated as mixed. A schematic representation of these possibilities is shown in Fig. S1.

To quantify this contribution, let *z* denote the average number of native contacts per residue, as introduced above, and define *Z* = 2*z* as the average number of bonds, or equivalently the average coordination number, per residue. For representative globular proteins, *z* is typically found to be close to 3, corresponding to *Z* ≈ 6, as illustrated in Fig. S2. In terms of this coordination measure, the total number of distinct shared-residue contact pairs in a chain of *N* residues is denoted by *W*_tot_. Since each residue with average coordination number *Z* can anchor 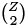 distinct pairs of incident contacts, the total number of such wedge motifs is

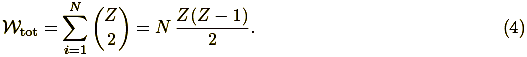

The quantity *W*_nn_ denotes the number of native–native wedges, namely those configurations in which the central residue is native and both residues connected to it within the wedge are also native. Within a mean-field treatment, the corresponding average number of such wedges is given below in 5; the detailed derivation of this reduction to the mean-field form is provided in Section S1 of the Supporting Information.

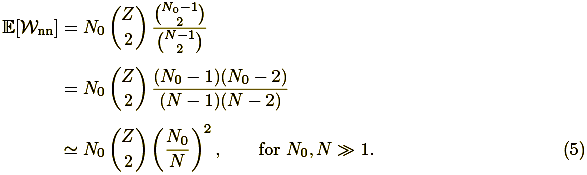

The remaining wedges are classified as mixed cooperative wedges, so that *W*_mix_ = *W*_tot_ − *W*_nn_.

### One-Dimensional Native-Cooperative Folding Model

The shared-residue cooperative interaction is incorporated into the basic one-dimensional spin-glass Hamiltonian through three additional parameters: 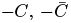, and Δ*C*. Here, −*C* denotes the cooperative stabilization energy associated with a native cooperative wedge, whereas 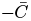 characterizes the corresponding non-native cooperative stabilization for a mixed wedge, with 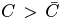 so that native cooperative stabilization is more pronounced than non-native cooperative stabilization. These two contributions enter the average energy together with the primary, secondary, and tertiary interaction strengths already present in Eq. (2). The parameter Δ*C* denotes the energetic variance associated with non-native cooperative stabilization and is correspondingly incorporated into the standard deviation of the one-dimensional spin-glass energy distribution in Eq. (3). With these cooperative contributions included, the corresponding mean energy and standard deviation take the following form:

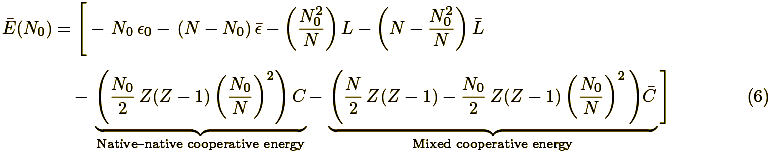

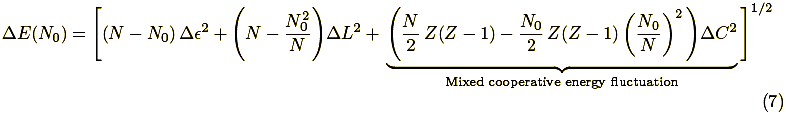

The thermodynamic description is formulated within the Gaussian energy approximation, in which configurations at fixed *N*_0_ are completely characterized by the mean *Ē*(*N*_0_) and the variance Δ*E*(*N*_0_)^2^. Within this framework, the density of states of the one-dimensional native cooperative folding model is given by

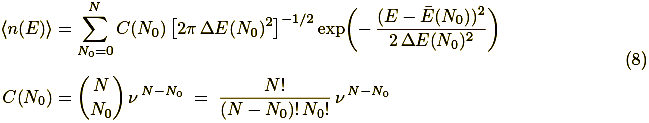

where *C*(*N*_0_) gives the number of configurations containing *N*_0_ native residues. The first line of Eq. (8) represents the sum over Gaussian energy distributions for all possible values of *N*_0_, each weighted by its corresponding multiplicity, while the second line counts the number of ways to choose the native residues together with the remaining non-native configurational degeneracy. Applying Boltzmann’s principle to Eq. (8), the microcanonical entropy in the thermodynamic limit takes the form

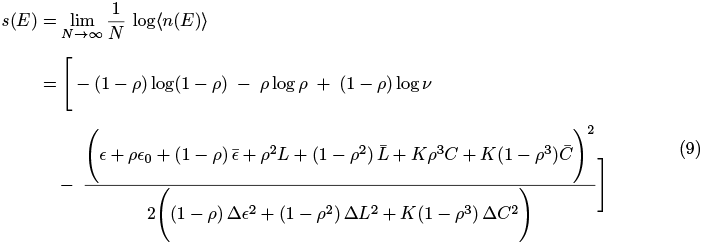

where 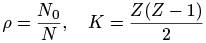.

Application of the thermodynamic identity 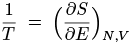 to the entropy in Eq. (9) yields the temperature-parametrised energy of the one-dimensional cooperative folding model:

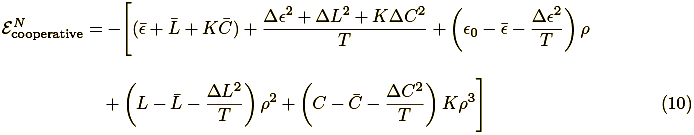

The corresponding canonical entropy and Helmholtz free energy of the one-dimensional cooperative folding model are given by

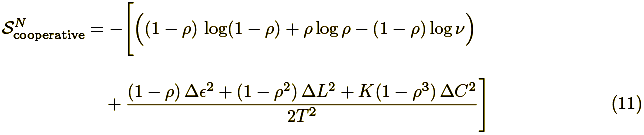

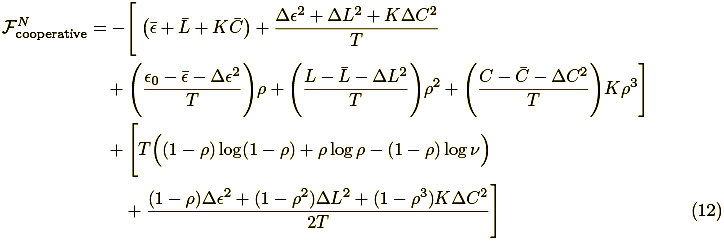

The phase behavior of the one-dimensional cooperative folding model is determined by minimizing the canonical free energy with respect to the native fraction *ρ*. The corresponding stationarity condition is

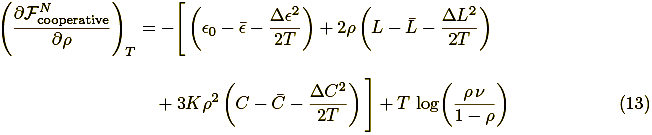

The equilibrium native fraction, *ρ*^⋆^, is obtained by solving Eq. (13) with 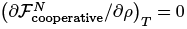, and provides the central order parameter describing the folding response under cooperative stabilization.

### Native-Wedge Folding Model with Weighted Cooperativity

A natural extension of the native-cooperative framework is to allow only a finite fraction of native– native wedges to contribute the full cooperative stabilization. This generalization is introduced to examine how much native cooperative bias is required to produce an effective folding scenario, rather than assuming the fully cooperative limit from the outset. To this end, we define a cooperativity fraction *f* ∈ [0, 1], which specifies the fraction of native–native wedges assigned cooperative stabilization. The limiting cases *f* = 0 and *f* = 1 correspond, respectively, to the absence of native cooperative bias and to the fully native-cooperative model. Within this generalized description, the modified mean energy and standard deviation are given by the following expressions:

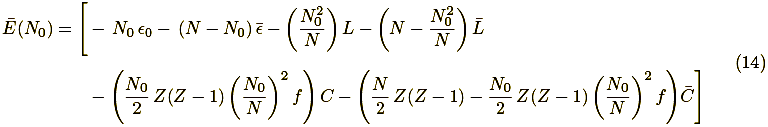

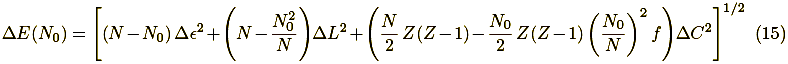

The corresponding density of states and microcanonical entropy may be constructed following the same procedure outlined in the earlier subsection for the native cooperative folding model. The derivative of the microcanonical entropy with respect to energy then provides the bridge to the canonical description through 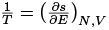, from which the temperature-dependent internal energy may be obtained. The corresponding canonical entropy and free energy then follow from the same formal procedure, and their detailed expressions are provided in Section S2 of the Supporting Information.

Construction of the phase diagram for the weighted native-cooperative folding model requires minimization of the canonical free energy with respect to the native fraction *ρ*. The corresponding stationarity condition is

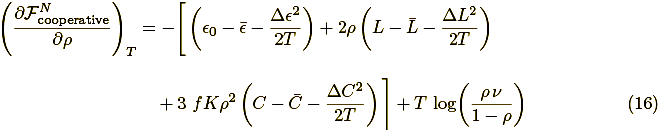

The equilibrium native fraction, *ρ*^⋆^, is obtained by setting Eq. (16) equal to zero and identifying the corresponding minimum of the free energy; it then serves as the order parameter used to determine the phase diagram of the weighted native-cooperative model.

## Results & Discussion

### Minimal Cooperative Stabilization Induces Folding in the One-Dimensional Spin-Glass Model

The equilibrium native fraction *ρ* is determined by solving the stationarity condition in Eq. (13). To enable a direct comparison with the earlier one-dimensional spin-glass model in the absence of cooperativity, we consider the baseline parameter set 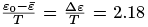 and 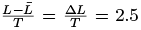. For these values, the noncooperative limit 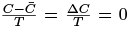 places the system in the unfolded regime, as indicated in the phase diagram of the original formulation by Wolynes and co-workers [14]; the same unfolded-state behavior for this parameter set has also been reported by Mitra *et al*. [42]. Against this reference point, the effect of cooperativity can be examined directly in terms of the native–non-native cooperative energy imbalance 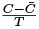 and the non-native cooperative energetic dispersion 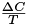.

The folding response is mapped systematically in the cooperative parameter space by varying 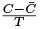 over the range [0, 5] in increments of 1, and 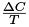 over the range [0, 2.5] in increments of 0.5, while fixing *Z* = 6, as discussed earlier. At each grid point, Eq. (13) is solved to obtain the equilibrium native fraction *ρ*, which serves as the order parameter for the phase diagram shown in Fig. 1(a). Within this description, the unfolded regime is associated with low native fractions, typically *ρ* ≈ 0.1–0.2, whereas the folded regime is associated with high native fractions, typically *ρ* ≈ 0.8–0.9.

**Figure 1.**
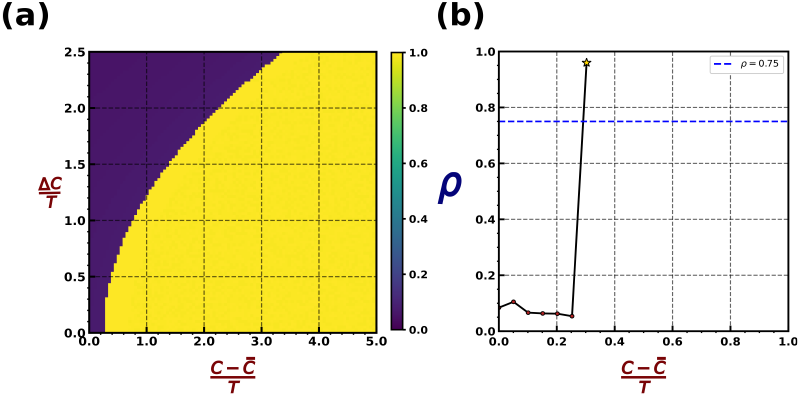
Cooperative stabilization drives the unfolded-to-folded transition in the one-dimensional native-cooperative model. (a) Phase diagram in the 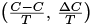 plane, constructed from the equilibrium native fraction *ρ*. (b) Folding onset profile showing the equilibrium native fraction *ρ* as a function of the native cooperative bias 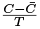 at fixed 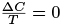. The onset of folding is identified by the crossing of the threshold *ρ* = 0.75, occurring at around 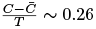.

Figure 1(a) displays the phase diagram in the 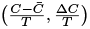 plane. The magenta region correspond to low native fractions, typically 0 < *ρ* < 0.2, and therefore signifies the unfolded regime, whereas the yellow region corresponds to high native fractions, with *ρ* 0.9, identifying the folded regime. Figure 1(b) shows the variation of the equilibrium native fraction *ρ* with the native cooperative bias 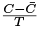 at fixed 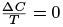. The onset of folding is identified by the crossing of the threshold value *ρ* = 0.75, in accordance with the folding criterion introduced by Wolynes and co-workers. The system undergoes a transition from the unfolded to the folded state at around 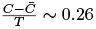. Taken together, these results show that the onset of folding occurs at a comparatively small native cooperative bias, beyond which the system is driven from the unfolded regime into a stable folded state.

For completeness, the consequence of using the same baseline parameter set as in the main analysis, namely 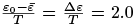 and 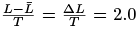, is presented separately in Fig. S3 of the Supporting Information. Under this equal parametrization, the primary and secondary–tertiary native stabilization contributions entering the first two terms of Eq. (13) aare effectively neutralized (as shown in the Supporting Information, Eq. ***26***), so that the folding transition is governed primarily by 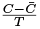 and 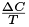 alone. This supplementary analysis leads to the same physical conclusion: even a comparatively small cooperative bias is sufficient to drive the system across the folding threshold and stabilize the folded state, even in the absence of any net stabilizing contribution from the primary and secondary–tertiary interactions.

### Parity of Native–Non-Native Deviation and Spread in Secondary–Tertiary Contact Stabilization Raises the Folding Threshold

A systematic examination of how secondary–tertiary native stabilization and its accompanying nonnative fluctuation influence cooperative folding is carried out by fixing the primary native energy gap at 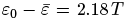, together with the corresponding primary energetic fluctuation Δ*ε* = 2.18 *T*, in keeping with the original spin-glass Hamiltonian parametrization, and taking *Z* = 6, as discussed earlier.The differential native–non-native secondary–tertiary stabilization, 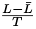, and the corresponding non-native secondary–tertiary energetic fluctuation, 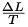, are then parametrized by imposing the equalvalue condition 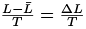, with values varied from 0.5 to 5.0 in steps of 0.5. For each such choice, the cooperative parameter space is mapped by varying 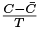 over the range [0, 5] and 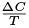 over the range [0, 2.5], following the same grid construction introduced in the previous subsection, and the optimal native fraction *ρ* is determined from the corresponding free-energy minimization.The resulting *ρ*-values are then used to construct the phase diagrams shown in Fig. 2. For clarity, only the representative cases 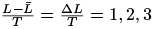 and 4 are displayed in the main text, while the remaining parameter-resolved phase diagrams are provided in Fig. S4 of the Supporting Information.

**Figure 2.**
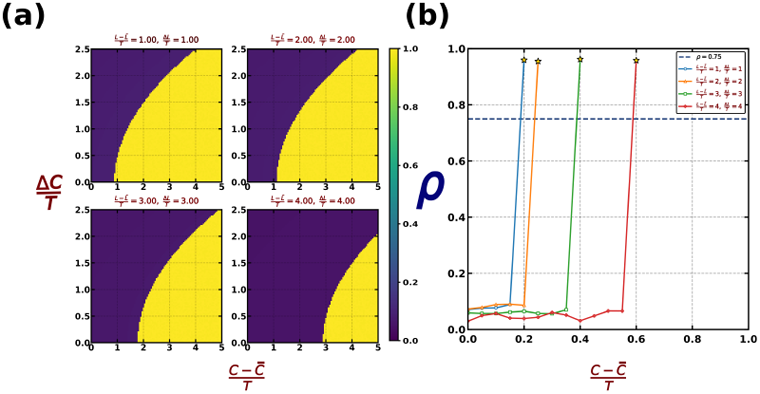
Secondary–tertiary native-stabilization–fluctuation parity alters the cooperative folding threshold. (a) Phase diagrams in the 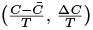 plane for representative matched values 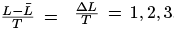, and 4. (b) Onset of the unfolded-to-folded transition, shown by the variation of the equilibrium native fraction *ρ* with 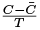 at fixed 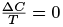.

Figure 2(a) shows that increasing the matched values of 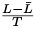 and 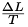 shifts the cooperative phase boundary progressively toward larger 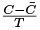, indicating that the onset of the folded regime requires an increasingly larger native cooperative bias. This shift is accompanied by a clear expansion of the unfolded region and a corresponding reduction of the folded region. The overall trend indicates that, although the increase in 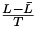 enhances the native–non-native secondary–tertiary stabilization, the simultaneous increase in 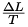 has a stronger destabilizing effect.

The same conclusion is reflected more directly in Fig. 2(b), which tracks the onset of folding at fixed 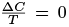. The minimum cooperative stabilization required to cross the folding threshold increases systematically with the matched choice 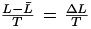: it is approximately 0.17 for 1, 0.20 for 2, 0.37 for 3, and 0.58 for 4. This trend shows that fluctuation in the secondary–tertiary interaction term affects the cooperative network more strongly than the corresponding mean native bias. As a result, even when 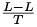 and 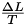 are increased by the same amount, the fluctuation term dominates and the cooperative threshold for folding shifts to higher values. Overall, the results indicate that the mean non-native secondary—tertiary stabilization and the spread in the corresponding interaction energies act in opposition, with the latter suppressing folding more strongly than the former promotes it.

### Baseline Native Bias–Fluctuation Balance Raises the Cooperative Folding Threshold

A complementary analysis is performed by keeping the secondary–tertiary contribution fixed at 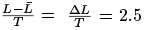, while varying the primary native–non-native energy difference and the corresponding primary energetic fluctuation under the equal-value condition 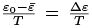. The representative values considered are 1.0, 1.5, 2.0, and 2.5. For each of these choices, the cooperative parameters 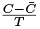 and 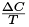 are varied over the same ranges used in Figs. 1 and 2, and the free energy is minimized to obtain the corresponding equilibrium values of *ρ*. These equilibrium native fractions are then used to construct the associated phase diagrams. Figure 3(a) shows a trend analogous to that observed for the equal secondary–tertiary mean–spread scenario discussed in Section. As the matched primary condition 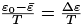 is increased, the folded region progressively contracts while the unfolded region expands. Figure 3(b) shows the variation of the equilibrium native fraction *ρ* with 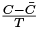 at fixed 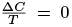 for the corresponding parameter sets. The threshold crossing shifts to larger 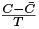 as the matched primary native-stabilization–fluctuation scale is increased, indicating that a higher cooperative stabilization is required to induce folding. Overall, the results show that under equal primary mean–spread parametrization, the fluctuation contribution dominates and shifts the cooperative folding threshold to higher values.

**Figure 3.**
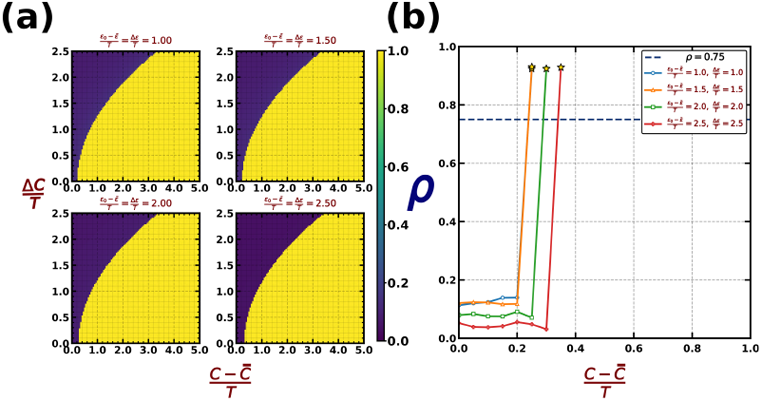
Primary native energy deviation–fluctuation parity shifts the cooperative folding boundary. (a) Phase diagrams in the 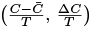 plane for representative matched choices 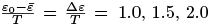, and 2.5. (b) Folding-onset curves showing the equilibrium native fraction *ρ* as a function of 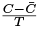 at fixed 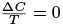.

A key quantity in this context is the free-energy barrier separating the unfolded and folded basins at coexistence. For the baseline one-dimensional spin-glass model, taking 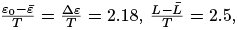, and 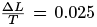 yields a double-well free-energy profile with two minima of equal depth, as shown in Supporting Fig. S5(a), thereby providing the appropriate reference point for estimating the folding barrier. Under these conditions, the barrier height is only Δ*F*^*‡*^ ≈ 0.035 *k*_B_*T*. In comparison, in the wedge-based cooperative model, we keep 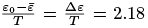 and 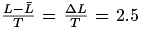 fixed, consistent with the parameter choice introduced earlier, and tune the cooperative interaction parameters to 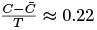 and 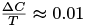. This yields an analogous coexistence point with equally stable unfolded and folded basins, as illustrated in Supporting Fig. S5(b), but now with a substantially larger barrier, Δ*F*^*‡*^ ≈ 0.45 *k*_B_*T*. The barrier is therefore enhanced by ~ 10-fold relative to that of the baseline one-dimensional spin-glass model. Physically, this increase reflects the fact that, once cooperative wedge motifs form along the folding pathway, the native state is stabilized not only by the underlying secondary–tertiary interactions but also by a mutually reinforcing cooperative network. In this sense, the present mean-field wedge model likely provides a lower-bound estimate of the barrier increase due to explicit cooperativity. For a protein of *N* residues, the number of possible cooperative residue-pair couplings can, in principle, scale up to *O*(*N* ^4^), providing an upper-limit estimate of the folding barrier attainable through extensive cooperative stabilization. Protein-folding experiments and molecular-dynamics simulations often suggest free-energy barriers in the 3–4 *k*_B_*T* range, suggesting that the true cooperative stabilization operative in protein folding lies between the present wedge-based mean-field estimate and the corresponding fully cooperative limit.

### Higher Fractional Cooperativity Lowers the Folding Threshold

The motivation for introducing fractional cooperative stabilization follows directly from the behavior discussed in Section and Fig. 1.In the preceding analysis, two distinct parameter regimes were considered. The first corresponds to the baseline one-dimensional spin-glass setting, with 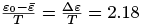 and 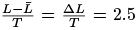, following the original noncooperative formulation; in this case, the system remains in the unfolded regime in the absence of any cooperative stabilization, with *ρ* confined to low values. The second corresponds to an equal-parametrization limit, 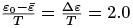 and 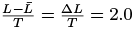, for which the contributions from these four baseline parameters effectively cancel in Eq. (13), thereby isolating a purely cooperativity-driven scenario. Despite their different physical interpretations, both regimes lead to the same qualitative outcome: the system crosses into the folded state at a comparatively small cooperative bias, of order 0.2–0.26. This immediately raises a natural question: how much of the total native residue-pair–residue-pair cooperative stabilization is actually required for the system to undergo folding, without invoking a uniformly strong cooperative network throughout the entire structure?

To address this point, we consider a partially cooperative native–native wedge model in which only a fraction *f* ∈ [0.1, 1.0] of the native residue-pair–residue-pair interactions is assigned the enhanced cooperative stabilization. In this construction, *f* = 0.1 corresponds to a situation in which only 10% of the native cooperative wedges are stabilized cooperatively, whereas *f* = 1.0 corresponds to the fully cooperative limit in which all such native wedges contribute the cooperative bias. The remaining fraction, 1 − *f*, of the native residue-pair–residue-pair sector is treated effectively as part of the nonnative cooperative background and is therefore associated with the weaker stabilization scale already represented by 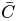. This parametrization allows us to examine, in a controlled manner, the minimal extent of cooperative coverage required for the system to cross into the folded regime, even when the net cooperative stabilization remains modest.

The equilibrium native fraction *ρ* is obtained by solving Eq. (16). In this analysis, the baseline primary and secondary–tertiary deviation and fluctuation parameters are all fixed at 2.0, with *Z* = 6, so that the folding response is governed predominantly by cooperative stabilization. The cooperative parameters 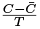 and 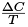 are varied over the same grid ranges used earlier for Figs. 1, 2, and 3. Figure 4(a) shows the unfolded–folded phase boundary in the 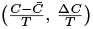 plane for fractional cooperative coverage *f* = 0.1–1.0. As *f* increases, the transition line shifts systematically toward lower 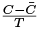, showing that a larger fraction of cooperative native wedges reduces the minimum cooperative bias required for folding. In particular, the threshold value of 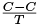 is approximately 1.8 for *f* = 0.1, decreases to about 0.9 for *f* = 0.2, ~ 0.6 for *f* = 0.3, ~ 0.45 for *f* = 0.4, and ~ 0.4 for *f* = 0.5. For *f* = 0.6–1.0, the transition lies within ~ 0.35 and decreases further to 0.2 in the fully cooperative limit, consistent with the behavior discussed earlier in Section.

**Figure 4.**
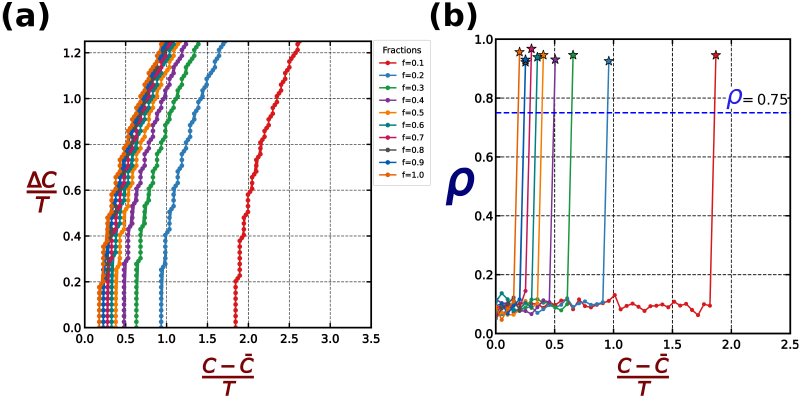
Incremental fractional native-wedge cooperativity lowers the folding threshold. (a) Unfolded– folded phase boundary in the 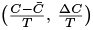 plane for cooperative coverage fractions *f* = 0.1–1.0. (b) Onset of the unfolded-to-folded transition, shown through the variation of the equilibrium native fraction *ρ* with 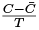 at fixed 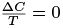 for the same set of *f* values.

Figure 4(b) shows the corresponding onset of folding through the variation of *ρ* with 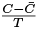 at fixed 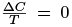 for different values of *f*. The unfolded-to-folded jump in *ρ* shifts to progressively lower 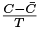 as the cooperative coverage is increased, in agreement with the phase-boundary trend seen in Fig. 4(a). Taken together, these results show that only a fraction of cooperative native wedges is sufficient to induce folding at a relatively small cooperative bias, whereas increasing cooperative coverage further lowers the folding threshold. This conclusion is also consistent with the distributional analysis reported by Best and co-workers [46] across several proteins, where the histogram projection of cooperative interaction pairs indicates that only a minority of native–native pairs, on the order of 20%–30%, are more strongly stabilized than the average over the combined native–nonnative and nonnative–nonnative pairs. From the mean-field perspective developed here, Figs. 4(a) and 4(b) show the same trend: even a modest degree of cooperative stabilization, of order 20%–30%, already corresponds to a minimal energetic bias of approximately 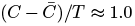, which is sufficient to drive the system across the folding threshold and stabilize the folded state. The full phase diagrams, including the complete *ρ*-resolved parameter space, are provided in Fig. S6 of the Supporting Information.

## Conclusion

Our study aims to show that introducing a shared-residue cooperative interaction into the one-dimensional spin-glass folding model qualitatively changes the folding landscape. Even in parameter regimes where the underlying one-dimensional model without cooperative stabilization remains unfolded, a comparatively small cooperative bias is sufficient to drive the system across the folding threshold and stabilize the folded state. This demonstrates that local cooperative residue-pair–residue-pair stabilization can act as an effective thermodynamic mechanism for promoting folding.

A second central result is that the dispersion in non-native interaction energies opposes folding more strongly than the corresponding mean native bias promotes it. This is seen both at the secondary– tertiary level and at the primary level: when the native–non-native deviation and the associated fluctuation are varied in equal increments, the unfolded region expands and the cooperative threshold shifts to higher values. Thus, disorder in the non-native sector weakens the effective cooperative drive and reduces foldability.

The partially weighted native-cooperative model further shows that full cooperative coverage is not required for folding. A finite fraction of cooperative native wedges is already sufficient to lower the folding threshold substantially, and the mean-field analysis indicates that even partial cooperative coverage, when accompanied by a modest bias toward native wedges, can drive the system from the unfolded to the folded regime. This indicates that folding need not rely on a uniformly cooperative network; instead, partial cooperative coverage can already provide the minimal stabilization required for folding.

A further important outcome is that cooperativity not only lowers the folding threshold but also increases the free-energy barrier separating the unfolded and folded basins. Relative to the corresponding one-dimensional spin-glass model, the wedge-based cooperative model yields a substantially larger coexistence barrier, indicating that cooperative wedge formation stabilizes folding through a mutually reinforcing network of native interactions.

Taken together, the present study highlights a simple physical picture: folding is controlled by the competition between cooperative stabilization and energetic fluctuation. Cooperative wedge interactions lower the folding threshold and also enhance the free-energy barrier protecting the folded state, whereas non-native fluctuation raises the threshold and suppresses foldability. From a biological perspective, this suggests that even modest local cooperative organization among native contacts may be sufficient to stabilize folding, while heterogeneity in non-native interactions can strongly oppose this process. In this sense, the model provides a minimal statistical-mechanical framework for understanding how local cooperativity reshapes the global folding landscape, while also highlighting a biologically relevant mechanism through which coordinated native interactions may promote efficient and robust folding in the presence of competing non-native disorder.

## Acknowledgments

The authors sincerely thank the Central Supercomputing Facility of the Indian Association for the Cultivation of Science, Kolkata and ANRF (ANRF grant ANRF/ARG/2025/003888/CS) for providing essential computational resources. Rohon Mitra gratefully acknowledges the Council for Scientific and Industrial Research (CSIR) for the award of a research fellowship, which supported this work.

---

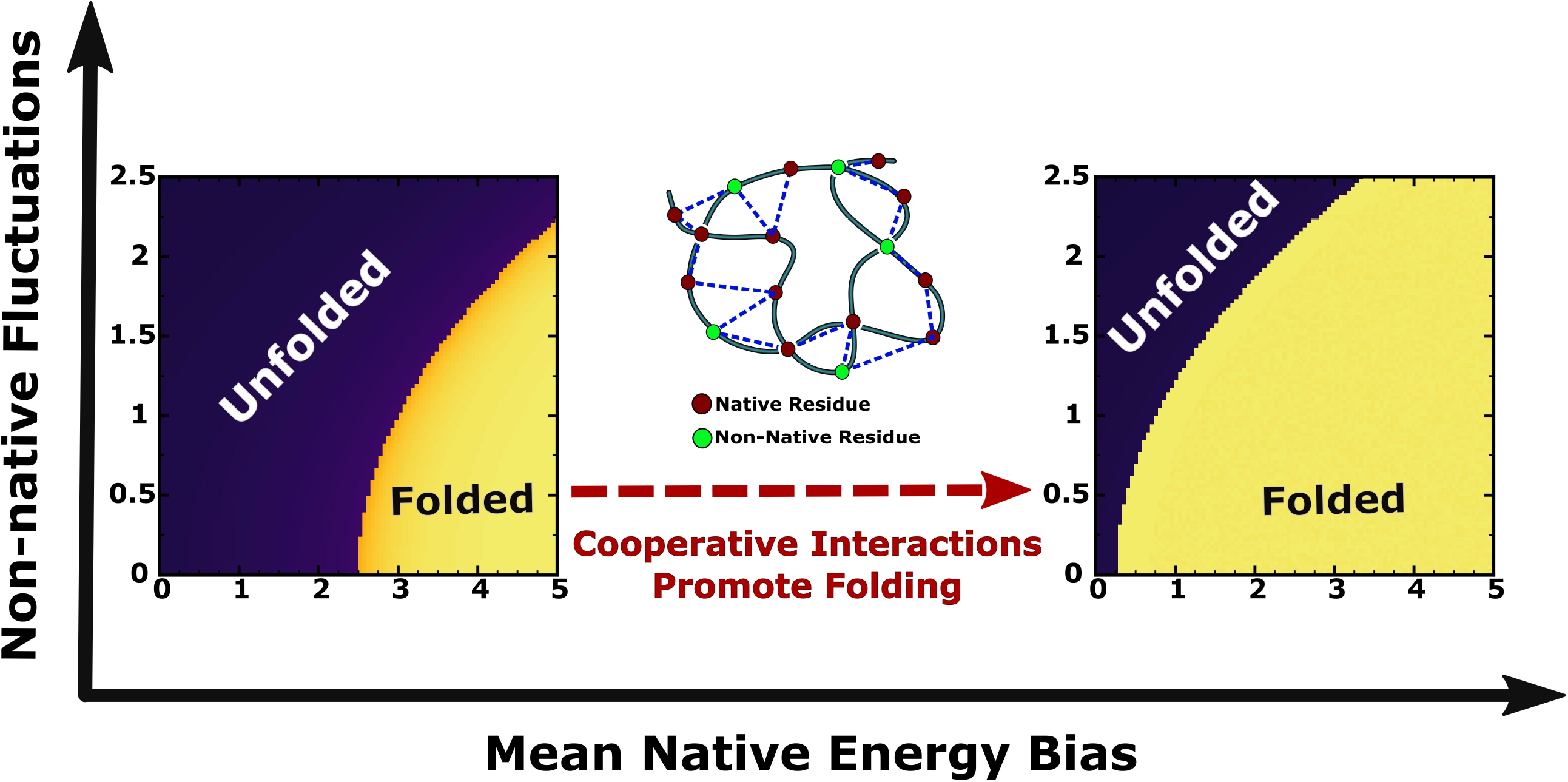

## References

[1] C. B. Anfinsen. The formation and stabilization of protein structure. Biochemical Journal, 128(4):737–749, 1972.

[2] Christian B. Anfinsen. Principles that govern the folding of protein chains. Science, 181(4096):223– 230, 1973.

[3] C. B. Anfinsen and H. A. Scheraga. Experimental and theoretical aspects of protein folding. Advances in Protein Chemistry, 29:205–300, 1975.

[4] Rainer Jaenicke. Protein folding: Local structures, domains, subunits, and assemblies. Biochemistry, 30(13):3147–3161, 1991.

[5] Roy R. Hantgan, Gordon G. Hammes, and Harold A. Scheraga. Pathways of folding of reduced bovine pancreatic ribonuclease. Biochemistry, 13(17):3421–3431, 1974.

[6] Mark Bycroft, Andreas Matouschek, James T. Kellis, Luis Serrano, and Alan R. Fersht. Detection and characterization of a folding intermediate in barnase by nmr. Nature, 346(6283):488–490, 1990.

[7] Sheena E. Radford, Christopher M. Dobson, and Philip A. Evans. The folding of hen lysozyme involves partially structured intermediates and multiple pathways. Nature, 358(6384):302–307, 1992.

[8] S. Walter Englander and Leland Mayne. The nature of protein folding pathways. Proceedings of the National Academy of Sciences of the United States of America, 111(45):15873–15880, 2014.

[9] Ken A. Dill and Hue Sun Chan. From levinthal to pathways to funnels. Nature Structural Biology, 4(1):10–19, 1997.

[10] Ken A. Dill. Polymer principles and protein folding. Protein Science, 8(6):1166–1180, 1999.

[11] Dmitry N. Ivankov and Alexei V. Finkelstein. Solution of levinthal’s paradox and a physical theory of protein folding times. Biomolecules, 10(2), 2020.

[12] Peter E. Leopold, Mauricio Montal, and José Nelson Onuchic. Protein folding funnels: A kinetic approach to the sequence-structure relationship. Proceedings of the National Academy of Sciences of the United States of America, 89(18):8721–8725, 1992.

[13] Natalya S. Bogatyreva and Alexei V. Finkelstein. Cunning simplicity of protein folding landscapes. Protein Engineering, 14(8):521–523, 2001.

[14] J. D. Bryngelson and P. G. Wolynes. Spin glasses and the statistical mechanics of protein folding. Proceedings of the National Academy of Sciences of the United States of America, 84(21):7524–7528, 1987.

[15] Joseph D. Bryngelson, José Nelson Onuchic, Nicholas D. Socci, and Peter G. Wolynes. Funnels, pathways, and the energy landscape of protein folding: a synthesis. Proteins, 21(3):167–195, 1995.

[16] José Nelson Onuchic, Zaida Luthey-Schulten, and Peter G. Wolynes. Theory of protein folding: the energy landscape perspective. Annual Review of Physical Chemistry, 48(1):545–600, 1997.

[17] José Nelson Onuchic and Peter G. Wolynes. Theory of protein folding. Current Opinion in Structural Biology, 14(1):70–75, 2004.

[18] Steven S. Plotkin and José N. Onuchic. Understanding protein folding with energy landscape theory part ii: Quantitative aspects. Quarterly Reviews of Biophysics, 35(3):205–286, 2002.

[19] Peter G. Wolynes, Jose N. Onuchic, and D. Thirumalai. Navigating the folding routes. Science, 267(5204):1619–1620, 1995.

[20] Peter G. Wolynes. Folding funnels and energy landscapes of larger proteins within the capillarity approximation. Proceedings of the National Academy of Sciences of the United States of America, 94(12):6170–6175, 1997.

[21] Joan Emma Shea and Charles L. Brooks. From folding theories to folding proteins: A review and assessment of simulation studies of protein folding and unfolding. Annual Review of Physical Chemistry, 52:499–535, 2001.

[22] Prem P. Chapagain, Jose L. Parra, Bernard S. Gerstman, and Yanxin Liu. Sampling of states for estimating the folding funnel entropy and energy landscape of a model alpha-helical hairpin peptide. Journal of Chemical Physics, 127(7), 2007.

[23] Hao Yu, Amar Nath Gupta, Xia Liu, Krishna Neupane, Angela M. Brigley, Iveta Sosova, and Michael T. Woodside. Energy landscape analysis of native folding of the prion protein yields the diffusion constant, transition path time, and rates. Proceedings of the National Academy of Sciences of the United States of America, 109(36):14452–14457, 2012.

[24] Kamalakar Gulukota and Peter G. Wolynes. Statistical mechanics of kinetic proofreading in protein folding in vivo. Proceedings of the National Academy of Sciences of the United States of America, 91(20):9292–9296, 1994.

[25] Kresten Lindorff-Larsen, Stefano Piana, Ron O. Dror, and David E. Shaw. How fast-folding proteins fold. Science, 334(6055):517–520, 2011.

[26] Robert L. Baldwin. The nature of protein folding pathways: The classical versus the new view. Journal of Biomolecular NMR, 5(2):103–109, 1995.

[27] Robert L. Baldwin and George D. Rose. Is protein folding hierarchic? ii. folding intermediates and transition states. Trends in Biochemical Sciences, 24(2):77–83, 1999.

[28] Eaton E. Lattman and George D. Rose. Protein folding — what’s the question? Proceedings of the National Academy of Sciences of the United States of America, 90(2):439–441, 1993.

[29] Zhuyan Guo and D. Thirumalai. Kinetics of protein folding: Nucleation mechanism, time scales, and pathways. Biopolymers, 36(1):83–102, 1995.

[30] Jean Francois St-Pierre, Normand Mousseau, and Philippe Derreumaux. The complex folding pathways of protein a suggest a multiple-funnelled energy landscape. Journal of Chemical Physics, 128(4), 2008.

[31] Hue Sun Chan, Zhuqing Zhang, Stefan Wallin, and Zhirong Liu. Cooperativity, local-nonlocal coupling, and nonnative interactions: Principles of protein folding from coarse-grained models. Annual Review of Physical Chemistry, 62:301–326, 2011.

[32] Andrew I. Jewett, Vijay S. Pande, and Kevin W. Plaxco. Cooperativity, smooth energy landscapes and the origins of topology-dependent protein folding rates. Journal of Molecular Biology, 326(1):247–253, 2003.

[33] Hüseyin Kaya and Hue Sun Chan. Solvation effects and driving forces for protein thermodynamic and kinetic cooperativity: How adequate is native-centric topological modeling? Journal of Molecular Biology, 326(3):911–931, 2003.

[34] Hue Sun Chan, Seishi Shimizu, and Hüseyin Kaya. Cooperativity principles in protein folding. Methods in Enzymology, 380:350–379, 2004.

[35] Robert B. Best, Gerhard Hummer, and William A. Eaton. Native contacts determine protein folding mechanisms in atomistic simulations. Proceedings of the National Academy of Sciences of the United States of America, 110(44):17874–17879, 2013.

[36] Victor Muñoz and William A. Eaton. A simple model for calculating the kinetics of protein folding from three-dimensional structures. Proceedings of the National Academy of Sciences of the United States of America, 96(20):11311–11316, 1999.

[37] Pierpaolo Bruscolini and Alessandro Pelizzola. Exact solution of the muñoz-eaton model for protein folding. Physical Review Letters, 88(25):258101, 2002.

[38] Wookyung Yu, Kwanghoon Chung, Mookyung Cheon, Muyoung Heo, Kyou Hoon Han, Sihyun Ham, and Iksoo Chang. Cooperative folding kinetics of bbl protein and peripheral subunit-binding domain homologues. Proceedings of the National Academy of Sciences of the United States of America, 105(7):2397–2402, 2008.

[39] Nicholas D. Socci and José Nelson Onuchic. Folding kinetics of protein like heteropolymers. Accepted in Journal of Chemical Physics, 1994.

[40] Leonid A. Mirny, Victor Abkevich, and Eugene I. Shakhnovich. Universality and diversity of the protein folding scenarios: A comprehensive analysis with the aid of a lattice model. Folding and Design, 1(2):103–116, 1996.

[41] N. D. Socci, J. N. Onuchic, and P. G. Wolynes. Diffusive dynamics of the reaction coordinate for protein folding funnels. Journal of Chemical Physics, 104(15):5860–5868, 1996.

[42] Rohon Mitra and Biman Jana. A model of protein folding with multiple native states: Metamor-phicity, intrinsic disorderness, and folding upon binding of proteins. Journal of Chemical Physics, 163(7):074110, 2025.

[43] Weihua Zheng, Nicholas P. Schafer, Aram Davtyan, Garegin A. Papoian, and Peter G. Wolynes. Predictive energy landscapes for protein-protein association. Proceedings of the National Academy of Sciences of the United States of America, 109(47):19244–19249, 2012.

[44] Ken A. Dill, Klaus M. Fiebig, and Hue Sun Chan. Cooperativity in protein-folding kinetics. Proceedings of the National Academy of Sciences of the United States of America, 90(5):1942– 1946, 1993.

[45] Zachary P. Gates, Michael C. Baxa, Wookyung Yu, Joshua A. Riback, Hui Li, Benoît Roux, Stephen B. H. Kent, and Tobin R. Sosnick. Perplexing cooperative folding and stability of a low-sequence complexity, polyproline 2 protein lacking a hydrophobic core. Proceedings of the National Academy of Sciences of the United States of America, 114(9):2241–2246, 2017.

[46] David Wang, Layne B. Frechette, and Robert B. Best. On the role of native contact cooperativity in protein folding. Proceedings of the National Academy of Sciences of the United States of America, 121(22):e2319249121, 2024.

[47] Zhuqing Zhang and Hue Sun Chan. Competition between native topology and nonnative interactions in simple and complex folding kinetics of natural and designed proteins. Proceedings of the National Academy of Sciences of the United States of America, 107(7):2920–2925, 2010.

[48] Arash Zarrine-Afsar, Stefan Wallin, A. Mirela Neculai, Philipp Neudecker, P. Lynne Howell, Alan R. Davidson, and Hue Sun Chan. Theoretical and experimental demonstration of the importance of specific nonnative interactions in protein folding. Proceedings of the National Academy of Sciences of the United States of America, 105(29):9999–10004, 2008.

[49] Tao Chen and Hue Sun Chan. Native contact density and nonnative hydrophobic effects in the folding of bacterial immunity proteins. PLoS Computational Biology, 11(5), 2015.

[50] Shilpa Yadahalli, V. V. Hemanth Giri Rao, and Shachi Gosavi. Modeling non-native interactions in designed proteins. Israel Journal of Chemistry, 54(8–9):1230–1240, 2014.

[51] Fernando Bruno Da Silva, Vinícius G. Contessoto, Vinícius M. De Oliveira, Jane Clarke, and Vitor B. P. Leite. Non-native cooperative interactions modulate protein folding rates. Journal of Physical Chemistry B, 122(48):10817–10824, 2018.

[52] Robert B. Best. How well does a funneled energy landscape capture the folding mechanism of spectrin domains? Journal of Physical Chemistry B, 117(42):13235–13244, 2013.

[53] Beth G. Wensley, Sarah Batey, Fleur A. C. Bone, Zheng Ming Chan, Nuala R. Tumelty, Annette Steward, Lee Gyan Kwa, Alessandro Borgia, and Jane Clarke. Experimental evidence for a frustrated energy landscape in a three-helix-bundle protein family. Nature, 463(7281):685–688, 2010.

[54] Beth G. Wensley, Lee Gyan Kwa, Sarah L. Shammas, Joseph M. Rogers, Stuart Browning, Ziqi Yang, and Jane Clarke. Separating the effects of internal friction and transition state energy to explain the slow, frustrated folding of spectrin domains. Proceedings of the National Academy of Sciences of the United States of America, 109(44):17795–17799, 2012.

[55] Feng Liu, Marcelo Nakaema, and Martin Gruebele. The transition state transit time of WW domain folding is controlled by energy landscape roughness. Journal of Chemical Physics, 131(19), 2009.

[56] Hoi Sung Chung, Stefano Piana-Agostinetti, David E. Shaw, and William A. Eaton. Structural origin of slow diffusion in protein folding. Science, 349(6255):1504–1510, 2015.

[57] Jae Yeol Kim, Fanjie Meng, Janghyun Yoo, and Hoi Sung Chung. Diffusion-limited association of disordered protein by non-native electrostatic interactions. Nature Communications, 9(1), 2018.

[58] Franco O. Tzul, Katrina L. Schweiker, and George I. Makhatadze. Modulation of folding energy landscape by charge-charge interactions: Linking experiments with computational modeling. Proceedings of the National Academy of Sciences of the United States of America, 112(3):E259–E266, 2015.

[59] Chi-Jui Feng, Ulrich Baxa, John M. Louis, and Hoi Sung Chung. Cooperative native contact formation facilitates free energy barrier crossing in protein folding. Physical Review Letters, 136(10):108401, 2026.

[60] Christopher A. Hunter and Harry L. Anderson. What is cooperativity? Angewandte Chemie International Edition, 48(41):7488–7499, 2009.

[61] A. Horovitz. The relation between cooperativity in ligand binding and intramolecular cooperativity in allosteric proteins. Proceedings of the Royal Society B: Biological Sciences, 259(1354):85–87, 1995.

[62] A. W. Lee and Martin Karplus. Structure-specific model of hemoglobin cooperativity. Proceedings of the National Academy of Sciences of the United States of America, 80(23):7055–7059, 1983.

[63] Gary K. Ackers, Michael L. Doyle, David Myers, and Margaret A. Daugherty. Molecular code for cooperativity in hemoglobin. Science, 255(5040):54–63, 1992.

[64] Yingwen Huang and Gary K. Ackers. Enthalpic and entropic components of cooperativity for the partially ligated intermediates of hemoglobin support a “symmetry rule” mechanism. Biochemistry, 34(19):6316–6327, 1995.

[65] Yingwen Huang and Gary K. Ackers. Transformation of cooperative free energies between ligation systems of hemoglobin: Resolution of the carbon monoxide binding intermediates. Biochemistry, 35(3):704–718, 1996.

[66] Jo M. Holt and Gary K. Ackers. Asymmetric distribution of cooperativity in the binding cascade of normal human hemoglobin. 2. stepwise cooperative free energy. Biochemistry, 44(36):11939–11949, 2005.

[67] Jo M. Holt, Alexandra L. Klinger, Connie S. Yarian, Varsha Keelara, and Gary K. Ackers. Asymmetric distribution of cooperativity in the binding cascade of normal human hemoglobin. 1. cooperative and noncooperative oxygen binding in zn-substituted hemoglobin. Biochemistry, 44(36):11925–11938, 2005.

[68] Gary K. Ackers and Jo M. Holt. Asymmetric cooperativity in a symmetric tetramer: Human hemoglobin. Journal of Biological Chemistry, 281(17):11441–11443, 2006.

[69] Jo M. Holt and Gary K. Ackers. The hill coefficient. inadequate resolution of cooperativity in human hemoglobin. Methods in Enzymology, 455:193–212, 2009.

[70] Jana L. Villemain and David P. Giedroc. Characterization of a cooperativity domain mutant lys3 → ala (k3a) t4 gene 32 protein. Journal of Biological Chemistry, 271(44):27623–27629, 1996.

[71] Lizhe Zhu, Daan Frenkel, and Peter G. Bolhuis. Role of fluctuations in ligand binding cooperativity of membrane receptors. Physical Review Letters, 106(16):168103, 2011.

[72] Jacob D. Marold, Thuy P. Dao, Tural Aksel, and Doug Barrick. Resolving cooperative interactions in protein folding. Biophysical Journal, 108(2):347a, 2015.

[73] Biman Jana, Dwaipayan Chakrabarti, and Biman Bagchi. Glassy orientational dynamics of rodlike molecules near the isotropic-nematic transition. Physical Review E, 76(1):011712, 2007.

[74] James L. Martin Robinson and Willem K. Kegel. Cooperative transitions involving hydrophobic polyelectrolytes. Proceedings of the National Academy of Sciences of the United States of America, 120(6):e2211088120, 2023.

[75] Elena Yu. Tupikina. Cooperativity of hydrogen bonds in biomolecular systems and their influence on structure and function. Coordination Chemistry Reviews, 549, 2026.

[76] Jakub Svoboda and Krishnendu Chatterjee. Promoters of cooperation in evolutionary games. Proceedings of the National Academy of Sciences of the United States of America, 122(51):e2524109122, 2025.

[77] Régis Turuban, Giovanni Noselli, Alfred Beran, and Antonio DeSimone. Cooperative mixing through hydrodynamic interactions in stylonychia lemnae. Proceedings of the National Academy of Sciences of the United States of America, 122(37):e2500588122, 2025.

[78] Jordi Piñero, Artemy Kolchinsky, Sidney Redner, and Ricard Solé. Neutral theory of cooperative dynamics. Proceedings of the National Academy of Sciences of the United States of America, 122(51):e2515423122, 2025.

