## Supplementary Information for "Local cooperative interactions reshape the folding transition in a one-dimensional spin-glass model"

### 1 Supplementary Equations

#### S1: Derivation of Shared-Residue ("Wedge") Cooperative Counts

In this section, we derive in detail the combinatorial expressions for the number of shared-residue cooperative units, which we refer to as *wedges*. A wedge is defined as an unordered pair of contacts sharing a common central residue. These objects provide a natural way to quantify local cooperative interactions in the single-state model.

Throughout this derivation, we assume that the contact network contains $N$ residues and that each residue has the same coordination number $z$. We denote by $X_{ij} \in \{0,1\}$ the contact indicator between residues $i$ and $j$, with

$$X_{ij} = X_{ji}, \qquad X_{ii} = 0,$$

and by $n_i \in \{0,1\}$ the indicator of whether residue $i$ is in the native state. The total number of native residues is therefore

$$\sum_{i=1}^{N} n_i = N_0. \tag{1}$$

A wedge anchored at residue $i$ is formed whenever residue $i$ is simultaneously connected to two distinct residues $j$ and $k$, with $j \neq k$. In other words, the pair of contacts $(ij, ik)$ constitutes one wedge centered at $i$.

##### S1.1 Total number of wedges

We first count the total number of wedges allowed purely by the topology of the contact graph. For a fixed residue $i$, there are exactly $z$ contacts incident on it because the degree of each residue is $z$. A wedge is obtained by choosing any two distinct contacts among these $z$ contacts. Hence the number of wedges centered at residue $i$ is

$$\mathcal{W}_i = \binom{z}{2} = \frac{z(z-1)}{2}. \tag{2}$$

Summing over all $N$ residues, the total number of wedges in the system is

$$\mathcal{W}_{\text{tot}} = \sum_{i=1}^{N} \mathcal{W}_i = \sum_{i=1}^{N} \binom{z}{2} = N\binom{z}{2} = N\,\frac{z(z-1)}{2}. \tag{3}$$

This is a purely geometric or topological quantity: it depends only on the contact connectivity and not on whether residues are native or non-native. The same count may also be written directly in terms of the contact matrix. For a fixed center $i$,

$$\mathcal{W}_i = \sum_{\substack{j<k \\ j\neq i,\, k\neq i}} X_{ij} X_{ik}. \tag{4}$$

Indeed, the product $X_{ij}X_{ik}$ equals unity only when both contacts $(i,j)$ and $(i,k)$ exist. Thus each unordered pair of neighbors of $i$ contributes exactly once, reproducing $\binom{z}{2}$.

#### S1.2 Exact number of native–native wedges

We now refine the counting by distinguishing wedges according to the native or non-native character of the participating residues. For a given residue $i$, define the number of its native neighbors by

$$d_i^N = \sum_{j \neq i} X_{ij} n_j. \tag{5}$$

This quantity counts how many neighbors of residue $i$ are native, since $X_{ij} = 1$ enforces the existence of a contact and $n_j = 1$ enforces that residue $j$ is native. A wedge centered at $i$ is called *native–native* if all three residues involved in the wedge are native: the central residue $i$ must satisfy $n_i = 1$, and both neighboring residues forming the two arms must also be native. If residue $i$ is native, then among its neighbors there are $d_i^N$ native ones, and the number of ways to choose two of them is

$$\binom{d_i^N}{2} = \frac{d_i^N(d_i^N - 1)}{2}.$$

If $i$ is not native, then no wedge centered at $i$ contributes to the native–native count. Therefore the exact number of native–native wedges is

$$\mathcal{W}_{\text{nn}} = \sum_{i=1}^{N} n_i \binom{d_i^N}{2} = \sum_{i=1}^{N} n_i \, \frac{d_i^N(d_i^N - 1)}{2}. \tag{6}$$

An equivalent explicit form may be obtained by expanding the binomial factor into a sum over unordered neighbor pairs:

$$\binom{d_i^N}{2} = \sum_{\substack{j<k \\ j \neq i,\, k \neq i}} X_{ij} X_{ik} n_j n_k. \tag{7}$$

Substituting this into Eq. (6) gives

$$\mathcal{W}_{\text{nn}} = \sum_{i=1}^{N} \sum_{\substack{j<k \\ j \neq i,\, k \neq i}} n_i \, X_{ij} X_{ik} \, n_j n_k. \tag{8}$$

This expression makes the counting transparent: $X_{ij} X_{ik}$ ensures that the two contacts exist and $n_i n_j n_k$ ensures that the central residue and both arm residues are all native.

#### S1.3 Definition of mixed wedges

All wedges that are not native–native are grouped into a complementary class, which we call *mixed wedges*. This class includes both wedges with a native centre but at least one non-native arm, and wedges whose centre itself is non-native. Since every wedge belongs either to the native–native class or to the mixed class, the number of mixed wedges is simply

$$\mathcal{W}_{\text{mix}} = \mathcal{W}_{\text{tot}} - \mathcal{W}_{\text{nn}}. \tag{9}$$

##### S1.4 Mean-field estimate under uniform mixing

We next derive a mean-field expression for the average number of native–native wedges under the assumption of uniform mixing. The idea is that, conditional on residue $i$ being native, its $z$ neighbors are chosen uniformly without replacement from the remaining $N-1$ residues. Among these $N-1$ residues, exactly $N_0-1$ are native.
Accordingly, the number $d_i^N$ of native neighbors of a native center $i$ follows a hypergeometric distribution:

$$d_i^N \,\big|\, (n_i = 1) \sim \text{Hypergeometric}(N-1,\ N_0-1,\ z). \tag{10}$$

Thus,

$$\Pr\big(d_i^N = m \,\big|\, n_i = 1\big) = \frac{\binom{N_0-1}{m}\binom{N-N_0}{z-m}}{\binom{N-1}{z}}, \qquad m = 0, 1, \ldots, z. \tag{11}$$

For a fixed native center $i$, the number of native–native wedges attached to it is $\binom{d_i^N}{2}$. Therefore,

$$\mathbb{E}\left[\binom{d_i^N}{2} \,\middle|\, n_i = 1\right] = \frac{1}{2}\,\mathbb{E}\big[d_i^N(d_i^N-1) \,\big|\, n_i = 1\big]\,. \tag{12}$$

Using the second falling-factorial moment of the hypergeometric distribution,

$$\mathbb{E}[(X)_2] = \frac{(n)_2(K)_2}{(M)_2},$$

with

$$M = N-1, \qquad K = N_0 - 1, \qquad n = z,$$

we obtain

$$\mathbb{E}\big[d_i^N(d_i^N-1) \,\big|\, n_i = 1\big] = \frac{(z)_2\,(N_0-1)_2}{(N-1)_2}. \tag{13}$$

Hence,

$$\mathbb{E}\left[\binom{d_i^N}{2} \,\middle|\, n_i = 1\right] = \frac{(z)_2\,(N_0-1)_2}{2\,(N-1)_2}. \tag{14}$$

Writing the falling factorials explicitly,

$$(z)_2 = z(z-1), \qquad (N_0-1)_2 = (N_0-1)(N_0-2), \qquad (N-1)_2 = (N-1)(N-2),$$

we obtain

$$\mathbb{E}\left[\binom{d_i^N}{2} \,\middle|\, n_i = 1\right] = \frac{z(z-1)}{2}\,\frac{(N_0-1)(N_0-2)}{(N-1)(N-2)}. \tag{15}$$

This may be written more compactly as

$$\mathbb{E}\left[\binom{d_i^N}{2} \,\middle|\, n_i = 1\right] = \binom{z}{2}\frac{\binom{N_0-1}{2}}{\binom{N-1}{2}}. \tag{16}$$

Finally, since there are $N_0$ native centers, the mean-field estimate for the total number of native–native wedges is

$$\mathbb{E}[\mathcal{W}_{\rm nn}] = N_0 \binom{z}{2} \frac{\binom{N_0-1}{2}}{\binom{N-1}{2}}. \tag{17}$$

Equivalently,

$$\mathbb{E}[\mathcal{W}_{\rm nn}] = N_0 \, \frac{z(z-1)}{2} \, \frac{(N_0-1)(N_0-2)}{(N-1)(N-2)}. \tag{18}$$

This result has a straightforward interpretation: it is the number of native centers, multiplied by the number of wedges per center, multiplied by the probability that both arms of a wedge connect to native residues.

##### S1.5 Summary of results

Collecting the expressions derived above, we obtain

$$\mathcal{W}_{\rm tot} = N\binom{z}{2} = N\,\frac{z(z-1)}{2}, \tag{19}$$

$$\mathcal{W}_{\rm nn} = \sum_{i=1}^{N} n_i \binom{d_i^N}{2} = \sum_{i=1}^{N} n_i \, \frac{d_i^N(d_i^N-1)}{2}, \tag{20}$$

$$\mathcal{W}_{\rm mix} = \mathcal{W}_{\rm tot} - \mathcal{W}_{\rm nn}, \tag{21}$$

$$\mathbb{E}[\mathcal{W}_{\rm nn}] = N_0 \binom{z}{2} \frac{\binom{N_0-1}{2}}{\binom{N-1}{2}} = N_0 \, \frac{z(z-1)}{2} \, \frac{(N_0-1)(N_0-2)}{(N-1)(N-2)}. \tag{22}$$

#### S2: Canonical Thermodynamic Functions for the Weighted Native-Wedge Cooperative Model

For the weighted native-wedge cooperative model, the canonical thermodynamic functions may be written as follows. The corresponding energy, entropy, and free-energy expressions are given by

$$\begin{aligned}\mathcal{E}^f_{\rm cooperative} = \Big[&(\bar{\epsilon} + \bar{L} + K\bar{C}) + \frac{\Delta\epsilon^2 + \Delta L^2 + K\Delta C^2}{T} \\ &+ (\epsilon_0 - \bar{\epsilon} - \tfrac{\Delta\epsilon^2}{T})\rho + \Big(L - \bar{L} - \tfrac{\Delta L^2}{T}\Big)\rho^2 + Kf\rho^3\Big(C - \bar{C} - \tfrac{\Delta C^2}{T}\Big)\Big]\end{aligned} \tag{23}$$

$$\begin{aligned}\mathcal{S}^f_{\rm cooperative} = &\Big[ -(1-\rho)\log(1-\rho) - \rho\log\rho + (1-\rho)\log\nu\Big] \\ &- \frac{(1-\rho)\Delta\epsilon^2 + (1-\rho^2)\Delta L^2 + K(1-f\rho^3)\Delta C^2}{2T^2}\end{aligned} \tag{24}$$

$$
\begin{aligned}
\mathcal{F}^{f}_{\text{cooperative}} = -\Bigg[ & (\bar{\epsilon} + \bar{L} + K\bar{C}) + \frac{\Delta\epsilon^2 + \Delta L^2 + K\Delta C^2}{2T} \\
& + (\epsilon_0 - \bar{\epsilon} - \tfrac{\Delta\epsilon^2}{2T})\rho + \left(L - \bar{L} - \tfrac{\Delta L^2}{2T}\right)\rho^2 + Kf\rho^3\left(C - \bar{C} - \tfrac{\Delta C^2}{2T}\right)\Bigg] \\
& + T\Big[\rho\log\rho + (1-\rho)\log(1-\rho) - (1-\rho)\log\nu\Big]
\end{aligned}
\tag{25}
$$

### 2 Supplementary Figures

#### Schematic Representation of Native and Mixed Cooperative Wedges

Local cooperative stabilization is modeled through an additional interaction associated with a shared-residue contact pair, or “wedge,” formed by two contacts anchored on the same central residue. With the central residue taken to be native, the wedge is classified by the states of the two partner residues. If both are native, the motif is a *native cooperative wedge*; if one or both are non-native, it is a *mixed cooperative wedge*. Here, $N$ denotes a native residue and $NN$ denotes a non-native residue. A schematic representation is shown in Fig. S1.

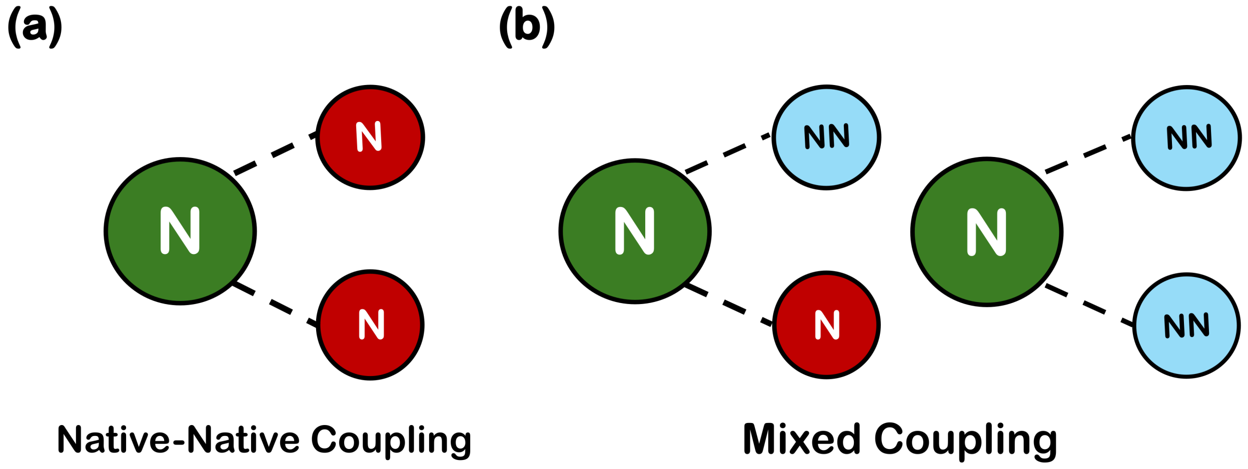

**Fig. S1:** Cooperative wedge motifs formed by two contacts sharing a common native central residue. Here, $N$ denotes a native residue and $NN$ a non-native residue. (a) Native–native coupling. (b) Mixed coupling, where one or both partner residues are non-native.

#### Estimation of the Average Coordination Number from Representative Protein Structures

A set of representative protein structures with PDB IDs 1amk, 1qgi, 1st3, 2ca2, 2jfn, 2xtp, and 5g4e was considered to estimate the characteristic local coordination in native protein folds. Using the shadow cut-off criterion applied to $C_\alpha$ atoms, we calculated the average number of native contacts per residue and the corresponding average number of bonds per residue, which may equivalently be viewed as the average coordination number. The resulting values are shown in Fig. S2 and provide a structural basis for the coordination parameters used in the cooperative wedge model.

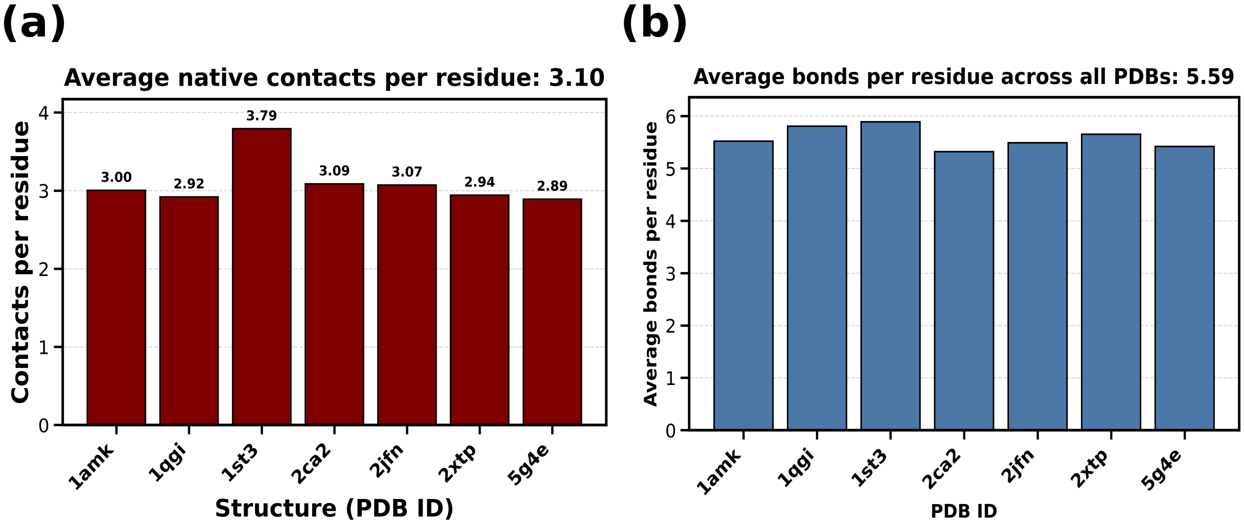

**Fig. S2:** Estimated native-contact coordination from representative globular protein structures. (a) Average number of native contacts per residue, $z$, for the selected PDB structures, giving an overall mean value of $z \approx 3.10$. (b) Corresponding average number of bonds per residue, $Z = 2z$, yielding an overall mean coordination value of $Z \approx 5.59$. These estimates support the use of $z \sim 3$ and $Z \sim 6$ as representative coordination measures in the wedge-based cooperative model.

#### Cooperative-Induced Folding Under Equal Baseline Parametrization

The role of cooperativity was further analyzed under the equal baseline parametrization

$$\begin{gathered}
\frac{\varepsilon_0 - \bar{\varepsilon}}{T} = \frac{\Delta\varepsilon}{T} = 2, \qquad \frac{L - \bar{L}}{T} = \frac{\Delta L}{T} = 2 \\
\varepsilon_0 - \bar{\varepsilon} = 2T, \qquad \Delta\varepsilon = 2T, \qquad L - \bar{L} = 2T, \qquad \Delta L = 2T \\
\varepsilon_0 - \bar{\varepsilon} - \frac{\Delta\varepsilon^2}{2T} = 2T - \frac{(2T)^2}{2T} = 2T - \frac{4T^2}{2T} = 2T - 2T = 0 \\
L - \bar{L} - \frac{\Delta L^2}{2T} = 2T - \frac{(2T)^2}{2T} = 2T - \frac{4T^2}{2T} = 2T - 2T = 0 \\
\text{Hence,} \left(\varepsilon_0 - \bar{\varepsilon} - \frac{\Delta\varepsilon^2}{2T}\right) = 0 \quad \text{and} \quad \left(L - \bar{L} - \frac{\Delta L^2}{2T}\right) = 0.
\end{gathered} \tag{26}$$

Under this choice, the primary and secondary–tertiary stabilization terms do not contribute any net native-state bias. The importance of Fig. S3 is therefore to show that the folding transition remains cooperative in origin: even in the absence of net stabilization from the noncooperative terms, a relatively small native cooperative bias is sufficient to drive the system across the folding threshold and stabilize the folded state. Thus, Fig. S3 serves as a control calculation demonstrating that the onset of folding is not an artifact of the baseline parametrization used in the main analysis.

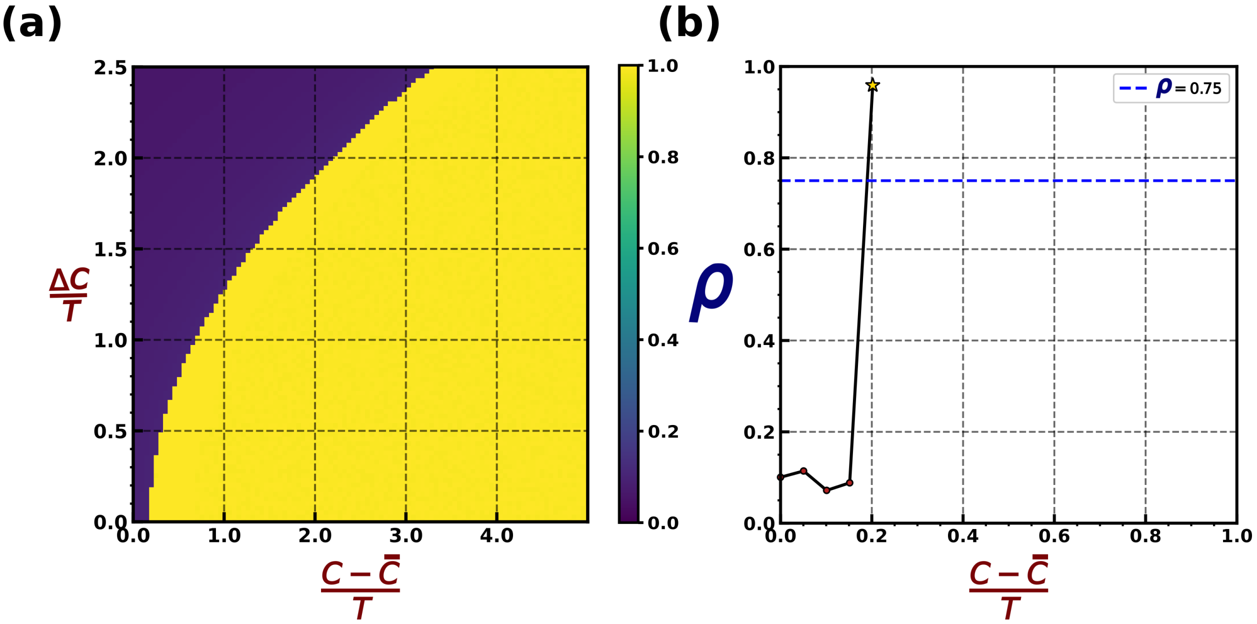

**Fig. S3:** Cooperative-induced folding under the equal baseline parametrization. (a) Phase diagram in the $\big((C-\bar{C})/T, \Delta C/T\big)$ plane constructed from the equilibrium optimized values of $\rho$. (b) Equilibrium native fraction $\rho$ as a function of $(C-\bar{C})/T$ at fixed $\Delta C/T = 0$, showing the onset of the unfolded-to-folded transition around $(C-\bar{C})/T \sim 0.2\,k_{\mathrm{B}}$.

#### Phase Diagrams for Matched Secondary–Tertiary Native–Non-Native Bias and Non-Native Fluctuation

The remaining parameter-resolved phase diagrams for the matched condition

$$\frac{L-\bar{L}}{T} = \frac{\Delta L}{T}$$

are shown in Fig. S4 for the values 0.5, 1.5, 2.5, 3.5, 4.5, and 5.0. These supplementary plots show the same trend as in the main text: as the equal secondary–tertiary bias and non-native fluctuation are increased together, the unfolded region expands and the folded region recedes. Correspondingly, the folding onset, or first-order transition boundary, shifts systematically toward higher values of $(C-\bar{C})/T$. This again indicates that the destabilizing effect of the non-native fluctuation term is stronger than the stabilizing effect of the corresponding mean native bias.

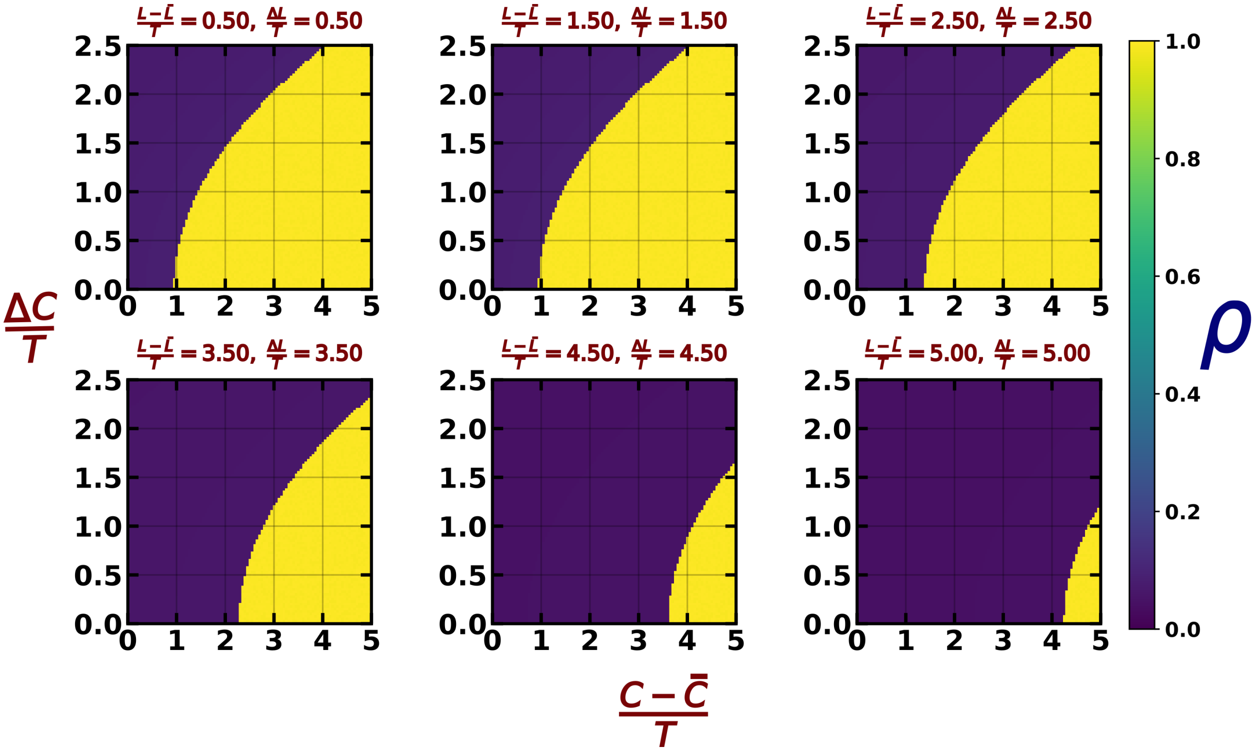

**Fig. S4:** Phase diagrams in the $\left(\frac{C-\bar{C}}{T}, \frac{\Delta C}{T}\right)$ plane showing the equilibrium optimized native fraction $\rho$. With increasing matched secondary–tertiary bias and non-native fluctuation, the phase boundary shifts to higher $\frac{C-\bar{C}}{T}$, indicating a higher folding onset.

#### Free-Energy Barrier Comparison Between the One-Dimensional Spin-Glass and Wedge-Based Cooperative Models

This analysis, shown in Fig. S5, compares the coexistence free-energy profiles of the one-dimensional spin-glass and wedge-based cooperative models, demonstrating that explicit cooperativity substantially increases the barrier separating the unfolded and folded basins.

#### Phase Diagrams for Fractional Native-Wedge Cooperativity

The full $\rho$-resolved phase diagrams for fractional native-wedge cooperativity, shown in Fig. S6, are presented here for $f = 0.1$ to 1.0. In each case, the equilibrium optimized native fraction $\rho$ is mapped in the $\left(\frac{C-\bar{C}}{T}, \frac{\Delta C}{T}\right)$ plane. The same trend discussed in the main text is retained throughout: as the fractional cooperative contribution $f$ increases, the unfolded–folded boundary shifts progressively toward lower $\frac{C-\bar{C}}{T}$. Thus, increasing $f$ reduces the minimum native cooperative bias required to induce folding.

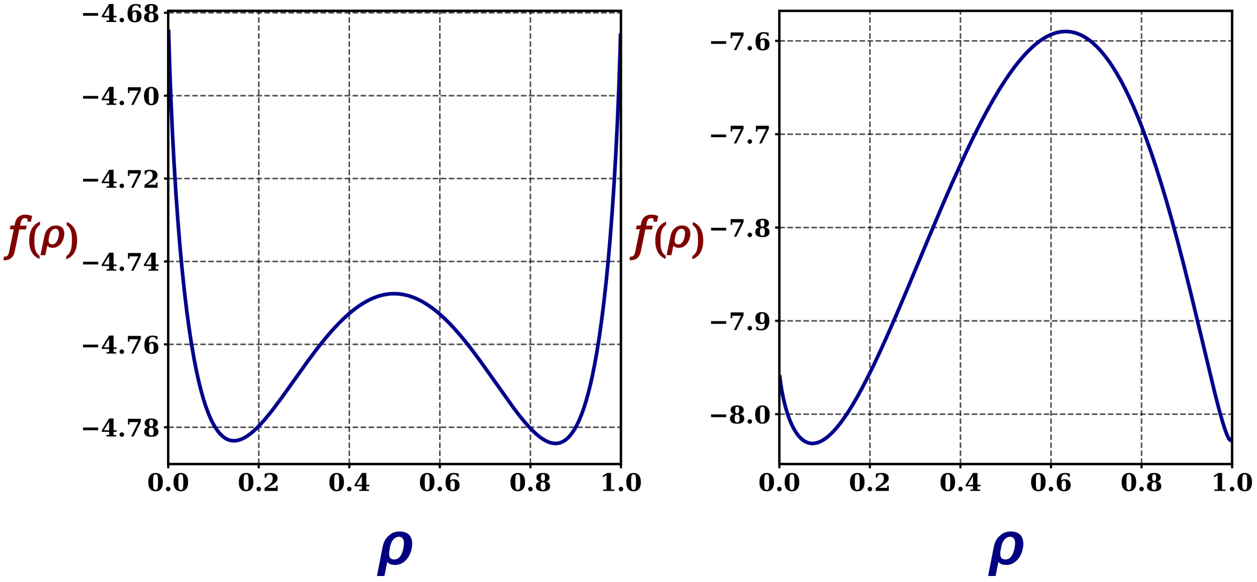

**Fig. S5:** Free-energy barrier comparison between the one-dimensional spin-glass model and the wedge-based cooperative model at coexistence. (a) Free-energy profile $f(\rho)$ of the one-dimensional spin-glass model, showing two equally stable basins separated by a shallow barrier, $\Delta F^{\ddagger} \approx 0.035\, k_{\mathrm{B}}T$. (b) Free-energy profile $f(\rho)$ of the wedge-based cooperative model, again showing coexistence of unfolded and folded basins but with a much larger barrier, $\Delta F^{\ddagger} \approx 0.45\, k_{\mathrm{B}}T$.

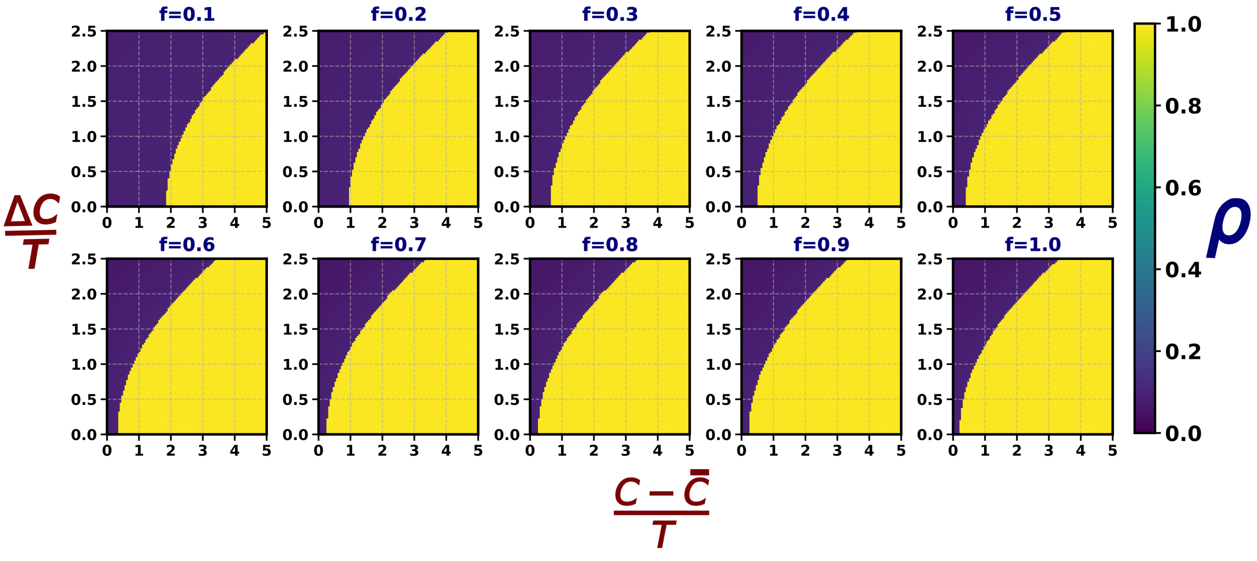

**Fig. S6:** Phase diagrams in the $\left(\frac{C-\bar{C}}{T}, \frac{\Delta C}{T}\right)$ plane for fractional native-wedge cooperativity $f = 0.1$ to 1.0, showing the equilibrium optimized native fraction $\rho$. With increasing $f$, the phase boundary shifts to lower $\frac{C-\bar{C}}{T}$, indicating a reduced folding threshold.
